# Beyond Bilingualism: multilingual experience correlates with caudate volume

**DOI:** 10.1101/209619

**Authors:** Alexis Hervais-Adelman, Natalia Egorova, Narly Golestani

## Abstract

The multilingual brain implements mechanisms that serve to select the appropriate language as a function of the communicative environment. Engaging these mechanisms on a regular basis appears to have consequences for brain structure and function. Studies have implicated the caudate nuclei as important nodes in polyglot language control processes, and have also shown structural differences in the caudate nuclei in bilingual compared to monolingual populations. However, the majority of published work has focused on the categorical differences between monolingual and bilingual individuals, and little is known about whether these findings extend to multilingual individuals, who have even greater language control demands. In the present paper, we present an analysis of the volume and morphology of the caudate nuclei, putamen, pallidum and thalami in 75 multilingual individuals who speak three or more languages. Volumetric analyses revealed a significant relationship between multilingual experience and right caudate volume, as well as a marginally-significant relationship with left caudate volume. Vertex-wise analyses revealed a significant enlargement of dorsal and anterior portions of the left caudate nucleus, known to have connectivity with executive brain regions, as a function of multilingual expertise. These results suggest that multilingual expertise might exercise a continuous impact on brain structure, and that as additional languages beyond a second are acquired, the additional demands for linguistic and cognitive control result in modifications to brain structures associated with language management processes.

## 1. Introduction

Multilingual individuals face an ongoing challenge in managing their language system. In order to efficiently communicate, a polyglot brain must implement mechanisms that permit the selection of the appropriate phonological, lexical and syntactic set for the current communicative environment, and the inhibition of the irrelevant ones. The mechanisms that allow language selection have been subject to investigation from multiple perspectives, which have yielded influential psycholinguistic models such as the Bilingual Interactivation + (Dijkstra & Van Heuven, 2002) and the Revised Hierarchical Model (Kroll, van Hell, Tokowicz, & Green, 2010), as well as comprehensive neurobiological accounts, such as the adaptive control hypothesis proposed by Green and Abutalebi (2013).

Brain imaging studies on bilingualism have largely revealed overlap between the brain functional language networks that are recruited during language processing in the first and second languages of bilinguals, with involvement of more heterogeneous regions in the L2 in less proficient bilinguals (Sebastian, Laird, & Kiran, 2011) and in late L2 learners (Berken, Gracco, & Klein, 2017; Indefrey, 2006), and with additional involvement of brain regions involved in language and executive control in these latter populations. With respect to language control, these studies have tended to support the view that language control and cognitive control processes depend upon similar networks. Regions associated with the executive control system, including the supplementary motor area and anterior cingulate cortex as well as the dorsal striatum are repeatedly implicated in tasks requiring language control (Abutalebi, 2008; Abutalebi et al., 2008; Abutalebi & Green*, 2008; Crinion et al., 2006; Hervais-Adelman, Moser-Mercer, & Golestani, 2011; Hervais-Adelman, Moser-Mercer, Michel, & Golestani, 2015). A mechanism whereby the basal ganglia may be implicated in polyglot language control is instantiated in the Conditional Routing model proposed by Stocco and collaborators (Stocco, Lebiere, & Anderson, 2010; Stocco, Yamasaki, Natalenko, & Prat, 2014), which posits that the basal ganglia coordinate cortico-cortical interactions when previously-learned cognitive routines cannot be applied, e.g. in cases of task-or language-switching. Findings of differences between mono- and bilinguals in brain regions involved in executive control can speak to reports of a “bilingual advantage” in various domains of cognitive function beyond language (Bialystok, 2011; Bialystok, Craik, & Luk, 2012; Diamond, 2010). The existence of such an advantage is disputed, and indeed effects are not always replicated (Paap & Greenberg, 2013; Paap, Johnson, & Sawi, 2015 2015). Nevertheless, we are minded to agree with Bialystok (2017) that it could be considered disingenuous to posit that the experience of multilingualism should have no effect on the brain at all. Indeed, there is substantial evidence, also from longitudinal studies, that bilingualism and language training influence brain function (Becker, Prat, & Stocco, 2016; Costumero, Rodriguez-Pujadas, Fuentes-Claramonte, & Avila, 2015), and also brain structure (Hervais-Adelman, Moser-Mercer, Murray, & Golestani, 2017; Stein et al., 2012). These findings are consistent with the notion that learning can indeed change the brain, even structurally, at micro-and macro-structural scales (Zatorre, Fields, & Johansen-Berg, 2012).

Structural imaging studies reveal divergent and heterogeneous findings regarding the brain structural differences between bilingual and monolingual individuals (García-Pentón, Fernández García, Costello, Duñabeitia, & Carreiras, 2015; Higby, Kim, & Obler, 2013; Luk & Pliatsikas, 2016). The earliest report of a reliable difference between bilingual and monolingual populations implicated the posterior supramarginal gyrus of the left inferior parietal lobule, which was found to exhibit a higher probability of more grey matter in bilingual than monolingual individuals (Mechelli et al., 2004). This grey matter difference showed a positive correlation with proficiency and a negative one with age of acquisition of the second language in the bilinguals. Since this report, numerous other brain areas have been shown to differ structurally between bilingual and monolingual individuals. In studies having used voxel-based morphometry (VBM), differences have been found in regions including, among others, cerebellum (Pliatsikas, Johnstone, & Marinis, 2014), left anterior temporal lobe (Abutalebi et al., 2014), anterior cingulate cortex (Abutalebi et al., 2015), left putamen (Abutalebi et al., 2013), Heschl’s Gyrus (Ressel et al., 2012), left caudate (Zou, Ding, Abutalebi, Shu, & Peng, 2012), bilateral caudate nuclei, putamen and thalamus (Burgaleta, Sanjuan, Ventura-Campos, Sebastian-Galles, & Avila, 2016). Effects of bilingualism have also been reported in white matter by several authors, in a variety of tracts, including the arcuate fasciculi (Hämäläinen, Sairanen, Leminen, & Lehtonen, 2017), the superior longitudinal fasciculi (Luk, Bialystok, Craik, & Grady, 2011; Pliatsikas, Moschopoulou, & Saddy, 2015), the inferior fronto-occipital fasciculi (Cummine & Boliek; Gold, Johnson, & Powell, 2013; Hämäläinen et al., 2017; Luk et al., 2011; Pliatsikas et al., 2015) and corpus callosum (Gold et al., 2013; Luk et al., 2011; Pliatsikas et al., 2015) and at a network level in fronto-temporal and fronto-parietal networks (Garcia-Penton, Perez Fernandez, Iturria-Medina, Gillon-Dowens, & Carreiras, 2014).

To date, only one previous study has compared cortical grey matter in multilingual individuals speaking more than two languages with bilinguals (Grogan et al., 2012). This investigation showed greater grey matter density in the right posterior supramarginal gyrus in the multilingual group than the bilingual group. One further study reports subcortical structural effects related to trilingualism - Abutalebi and colleagues (2013) showed an effect of language proficiency in the third language of trilingual individuals, such that the grey matter density of the left putamen increased as a function of proficiency. The diversity in results may arise from differences across studies in one or several out of a large number of confounding variables that also differentiate groups, other than language knowledge *per se*. These factors include immigrant status, cultural factors and socio-economic status, as well as how multiple languages are deployed (as considered by Green and Abutalebi, 2013 in the adaptive control hypothesis) and the context of acquisition (discussed by Pliatsikas, DeLuca, Moschoupoulou & Saddy, 2016). All of these factors have also been raised as potential confounding variables for the findings or lack thereof of a “bilingual advantage” (for a thorough overview of the controversy, see Paap et al., 2015; and rebuttals by Woumans & Duyck, 2015 and Bak, 2016).

In the present study, we aimed to overcome some of the potential confounds by exploring relationships between individual differences in brain structure in relation to multilingual expertise within a group of polyglot individuals, who mastered a minimum of three languages. Although the reasons for any given individual developing multilingual expertise may well be different, stemming from environmental, familial, motivational or educational factors, by focusing on an already multilingual population, systematic population-level confounds are less likely to influence findings. Furthermore, this approach allows us to explore brain structure in relation to language experience beyond bilingualism, and to reveal continuous relationships between multilingual experience and brain structure that are more easily attributable to language experience *per se* than might be provided by similar categorical comparisons between mono- and bi-lingual populations. We expected that structures most crucially implicated in the control and manipulation of multiple languages would be those most affected by multilingualism. We based our predictions on the results of a previous study of “extreme language control” (Hervais-Adelman et al., 2015), which implicated the caudate nucleus and the putamen in different cognitive levels of language control - the caudate in overarching task-level control and the putamen at a cognitively lower, more motoric, level of control. Here, we predicted that multilingual language experience beyond bilingualism, i.e. in individuals who speak 3 languages or more, would be systematically and positively related to the volumes of these two subcortical structures. It is worth noting that two previous studies (Burgaleta et al., 2016; Pliatsikas, DeLuca, Moschopoulou, & Saddy, 2016) have reported that the caudate nuclei of bilinguals are relatively larger compared to those of monolinguals, although Pliatsikas and colleagues found this only for highly proficient bilinguals who had not been highly immersed in an L2 environment. These studies also examined other subcortical structures and both found bilingualism-related structural changes in globus pallidus (bilaterally in the case of Pliatsikas et al., right only in the case of Burgaleta et al.) and thalamus (bilaterally in the case of Burgaleta et al. and right in the case of Pliatsikas et al.), and for comparability with these, we also analysed the globus pallidus and thalamus.

## 2. Methods

### 2.1 Participants and behavioral measures

Data were acquired in accordance with the Declaration of Helsinki, and with approval of the research ethics committees of the Lausanne and Geneva University Hospitals. Seventy-five individuals participated in the study (mean age: 25 years 11 months, s.d. 4 years 10 months, 42 female), all had completed or were engaged in at least tertiary education. They self-reported speaking three or more languages (range: 3-9, mean 4.37, s.d. 1.23), and were interviewed on their age of language acquisition (AoA) and proficiency levels in each of their reported languages. Weighted sums of AoA (earlier receiving higher weight) and proficiency (more proficient receiving higher weight) were calculated (cf. Hervais-Adelman, et al. 2015^1^) across languages spoken, to yield a compound and continuous index of language experience and proficiency (hereafter referred to as ‘LEXP’, mean: 35.15, s.d. 8.46). LEXP can be considered an aggregate measure of multilingual experience, by accounting for the contributions age of acquisition and language proficiency in addition to the total number of languages. This metric was employed in order to attempt to deal with the difficulty in establishing a summary measure that can incorporate multiple AoAs and proficiencies in a manner that can be comparable across participants having differing numbers of languages.

Sixty-seven of the datasets were acquired as part of a separate study (Hervais-Adelman et al., 2017), in which no analyses of the relationship between subcortical morphology and LEXP were carried out^2^. Of the 75 participants 40 had a acquired at least one second language before six years of age, and may be considered “early bilinguals”, and 35 only began acquiring their second and further languages after this age, and may be considered “late bilinguals”; these groups did not differ in terms of LEXP (overall: t(69.13)=0.385, p=.70, proficiency only: t(68.87)=1.34, p=0.19) or age (t(66.99)=1.03, p=.31).

### 2.2 Structural MRI

T1 MPRAGE images were acquired on the same model of scanner at two different sites, this being a Siemens 3T Trio MRI scanner, with an 8-channel head-coil (sagittal orientation, FoV: 240*256, slice thickness 1.2mm, 1mm * 1mm in-plane resolution, TR 2400ms, TE 2.98ms, Phase Encoding steps: 239, Flip angle 9°). Forty-eight participants were scanned at the Brain and Behaviour Laboratory, University of Geneva, and 27 at Lausanne University Medical Centre.

### 2.3 Subcortical Structure Extraction and Analysis

Subcortical structures of individual brains were extracted using FIRST (Patenaude, Smith, Kennedy, & Jenkinson, 2011), a utility supplied with FSL (Jenkinson, Beckmann, Behrens, Woolrich, & Smith, 2012). Segmentations of the left and right caudate nuclei, putamen, globus pallidus and thalamus were visually inspected for accuracy. In order to be able to account for the potential impact of head size on the volume of structures, estimated total intracranial volume (eTIV) was extracted, using the CAT12 toolbox in SPM12.

#### 2.3.1 Volumetric Analysis

All analyses were carried out in R (R Core Team). Initially, for each selected structure, weighted stepwise regression was employed to determine which covariates (from the following: Age, Sex, Handedness, eTIV, Scanner) should be included in an analysis of the contribution of LEXP to structural volume, using the WLE package (Agostinelli & Library, 2015). For all four structures under investigation (left caudate, right caudate, left putamen, right putamen), this analysis retained Age and eTIV as significant predictors of volume (Sex, Handedness and Scanner were therefore dropped from subsequent analyses). Robust regression analyses were executed using the “robust” package (Wang et al., 2014), including LEXP, Age and eTIV.

#### 2.3.2 Shape Analysis

The structures of interest were also submitted to a vertexwise analysis, in order to explore potential systematic differences in shape in relation to LEXP. Following the standard procedure implemented in FIRST, each structure was linearly registered (using 6 degrees of freedom) to the sample-specific average surface, mapped in MNI space. For each participant, a map was generated that contained the perpendicular vertexwise displacement vector required to map each vertex onto the mean. These values were then analysed using permutation-based non-parametric testing with Randomise (Stein, Winkler, Kaiser, & Dierks, 2014), and corrected for multiple comparisons using threshold free cluster enhancement (TFCE, Smith & Nichols, 2009). The design matrix contained the factor of interest (LEXP) and covariates of Age and eTIV (those retained by weighted stepwise regression as having explanatory power for the volumes of the structures of interest).

## 3. Results

Volumetric analyses revealed significant and marginally significant positive relationships between LEXP and right (t(71)=2.19, *p* = .032) and left caudate volumes (t(71) = 1.99, *p* = .050), respectively. These results are illustrated in Figure 1. Further examination showed that the Proficiency component of the LEXP measure is, at least qualitatively, more highly correlated with caudate volume (right: t(71)=2.24, *p* = .028; left: t(71)=2.02, *p* = .048) than is the AoA component (right: t(71)=1.81, *p* = .074; left: t(71)=1.85, *p* = .069). This result is consistent with the notion that linguistic expertise, more than the age of acquisition of a second language, is related to caudate volume, but is to be interpreted with a note of caution since the AoA and proficiency metrics are not independent of each other (as they are both dependent upon the number of languages reported by the participants). No relationship was found between putaminal volumes and LEXP (both left and right *p* > .75), nor between globus pallidus volumes and LEXP (left and right *p* > .19), nor between thalamus and LEXP (left and right *p* > .24).

A potential confounding issue is that age of acquisition has been shown to have an effect on the impact of second language acquisition on brain structure (cf. Mechelli et al., 2004; Kaiser et al., 2015). This is an intriguing issue given that languages learned earlier may be acquired implicitly, in contrast to those acquired later which may be learned explicitly, with consequences for the brain networks recruited (Morgan-Short, Steinhauer, Sanz, & Ullman, 2012). Unfortunately, the data we present here cannot fully address this specific question. The relatively weaker relationship between caudate volume and the AoA subcomponent of the LEXP metric, in comparison to that with the Proficiency component, suggests that it is cumulative proficiency, rather than precocity of acquisition, that is more closely related to caudate volume. Nevertheless, we carried out a supplementary analysis, using robust regression to evaluate whether those participants who acquired their first second language early showed different caudate volumes compared to those who acquired their first second language later, while controlling for the proficiency component of their LEXP score (eTIV and age were also included in the model, as before). These analyses revealed no main effect of early versus late bilingualism on caudate volume (left: p=0.11^3^, right: p=0.48), but there was a significant relationship between Proficiency and volume (left: t(71) = 4.90, *p* < .001, right: t(71) = 2.96, *p* = .004). Furthermore, there was no significant interaction between proficiency and early vs late bilingualism (left: p=0.158, right: p=0.63). These results suggest that there is no categorical distinction between early and late bilinguals in terms of the relationship between cumulative multilingual proficiency and caudate volume. While this is a relatively abstract datum, it is suggestive of the possibility that proficiency in multiple languages is indeed related to caudate volume. This does not, however, resolve the pressing question of causality in this relationship. Moreover, it is worth considering that with a group of individuals who speak a minimum of three languages, such an effect may be different than for bilinguals, and it is currently unknown whether any observed structural effect of early vs late bilingualism is mitigated or potentiated by subsequently-acquired languages.

**Figure 1:**
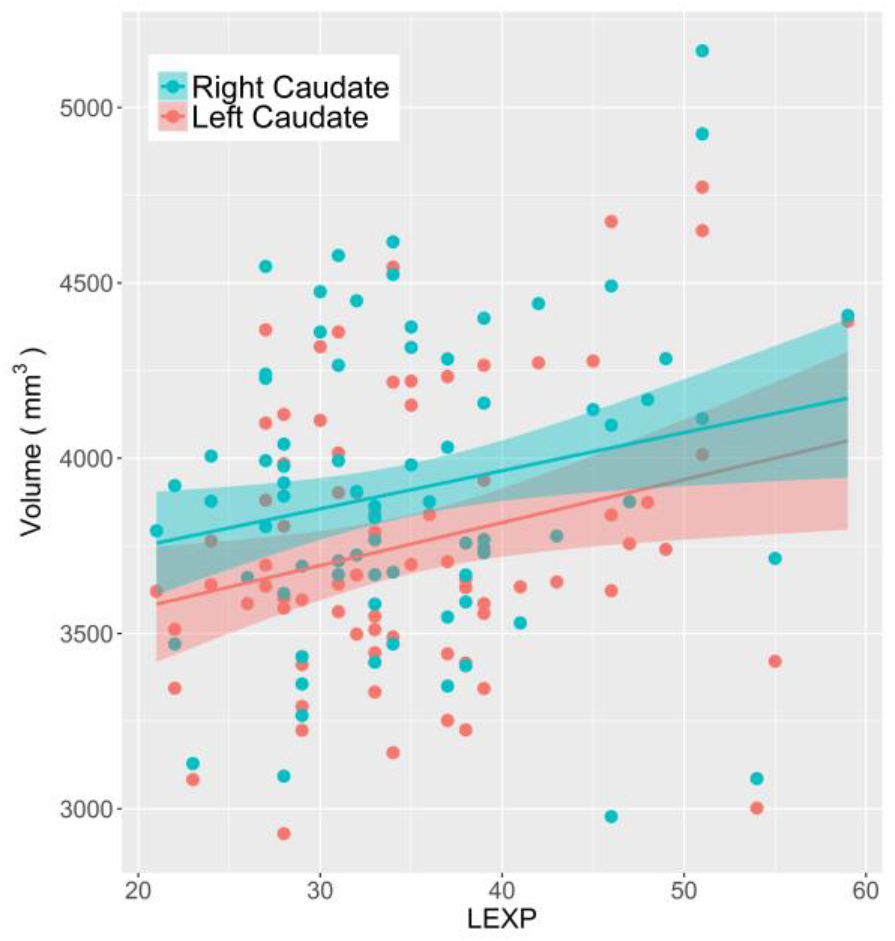
Scatter plot showing relationship between left and right caudate volumes and LEXP. Ribbons show 95% C.I. of robust linear regression.

Surface-based shape analysis of the structures revealed significant foci of expansion as a function of LEXP in two distinct clusters in the left caudate nucleus, one anterior and one dorso-medial (Figure 2). The likelihood of connectivity between these two caudate clusters and other brain regions was evaluated using the probabilistic Oxford-Imanova Striatal Connectivity Atlas with 7 sub-regions supplied with FSL. The anterior cluster (centre of mass, MNI co-ordinates, mm: −12, 24, −4) was assigned 58% likelihood of connectivity to the “executive” cortex, and the dorso-medial cluster (centre of mass, MNI co-ordinates, mm: −17, 3, 25) was assigned 31% likelihood of connectivity to executive cortex and 15% to caudal motor regions. No relationships between putaminal, pallidal or thalamic morphology and LEXP were revealed.

**Figure 2:**
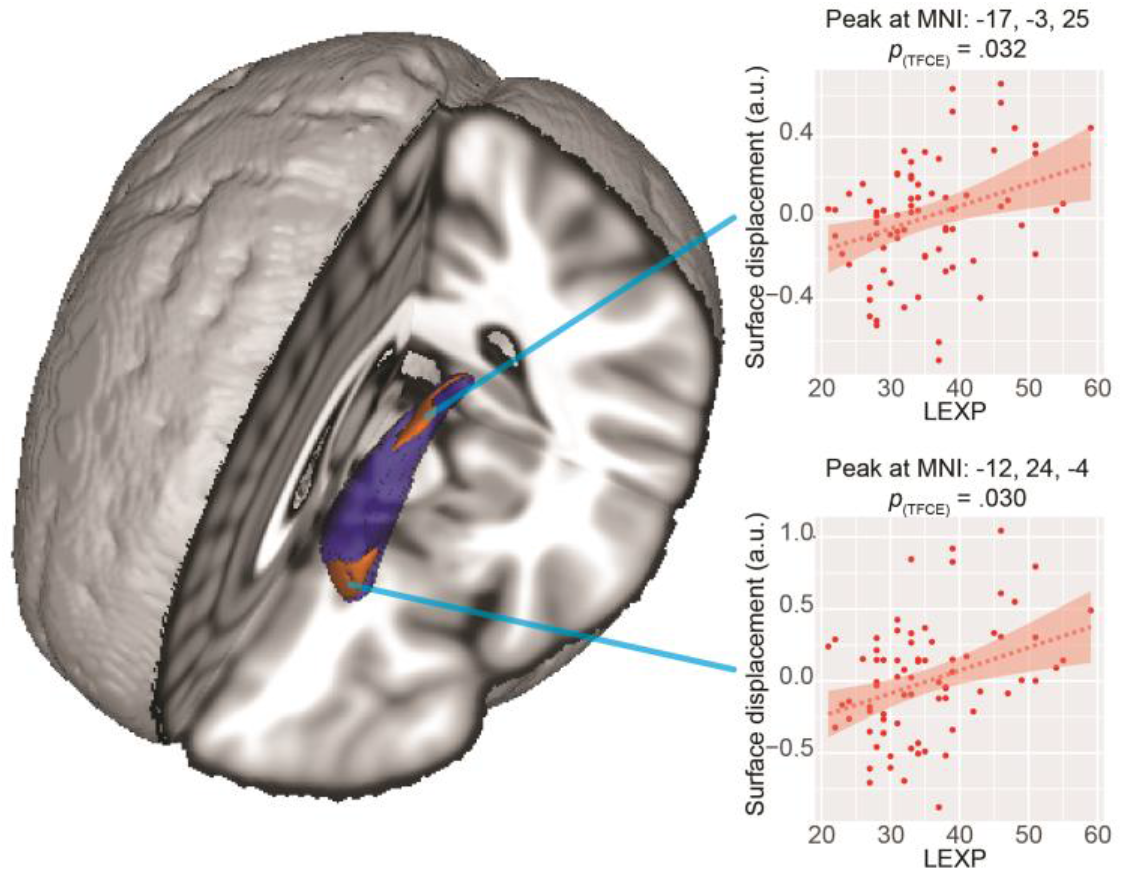
Rendering of standard MNI152 brain, highlighting left caudate (in blue), showing location of significant (p_(TFCE)_ < .05) outward surface displacement as a function of LEXP. Scatter plots illustrate displacement by LEXP at peak voxels of each indicated cluster. For illustration purposes only, trendlines show estimated robust linear regression, ribbons show 95% C.I.s for robust regression estimates.

## 4. Discussion

We find that increasing multilingual expertise correlates not only with bilateral caudate volume, but also with regionally-specific morphological alterations of the left caudate nucleus. These results support the view that polyglot individuals show structural adaptations that can be explained by their multilingual experience. To the best of our knowledge, this is the first demonstration of a continuous impact of increasing degrees of multilingualism on brain structure. By moving beyond the dichotomous comparison of monolingual with bilingual participants, we are able to more confidently put forward the view that the challenges of acquiring, maintaining and deploying multiple languages result in structural adaptation of the caudate nuclei.

The caudate nuclei have been shown to play a role in both language control (Crinion et al., 2006) and cognitive control (Grahn, Parkinson, & Owen, 2008; Hervais-Adelman et al., 2015), and have previously been shown to be enlarged in bilinguals vs. monolinguals (Burgaleta et al., 2016; Pliatsikas, DeLuca, Moschopoulou, & Saddy, 2016). Although there is also evidence for a role of the putamen in multilingual control, it may be that the absence of an observed relationship between putaminal structure and LEXP is due to the nature of its role: if, as suggested by Hervais-Adelman and colleagues (2015), the caudate is implicated in managing lexico-semantic sets as a function of task demands (c.f. the adaptive control hypothesis, Green & Abutalebi, 2013) while the putamen is involved in moment-to-moment suppression enabling the use of the appropriate language, it is conceivable that the number of competing languages does not substantially change the demands on this lower level of control. Such an interpretation is consistent with the cognitive roles of the caudate and putamen distinguished by Grahn and colleagues (Grahn et al., 2008), whereby the caudate has a role in the regulation of cognitive processes while the putamen is more central to movement control.

We cannot, based on the limited LEXP metric, distinguish between any potentially differential impact of proficiency, age of acquisition, factors relating to the process of acquiring, maintaining or storing multiple languages, or of more dynamic factors such as the context of language use and switching. Future work should strive to resolve this by acquiring more precise data on these factors in polyglot populations. Moreover, the LEXP score is derived from participants’ self-assessment, and as such is not a truly objective measure. We also note that our analysis cannot resolve the direction of causality in this relationship. Alternative explanations exist, which can only conclusively be addressed with longitudinal investigations. It is possible that individuals with relatively larger caudate nuclei have a particular aptitude for acquiring foreign languages, either through a cognitive advantage or as the result of motivational factors. It has been shown, for example, that caudate volumes positively correlate with IQ (Grazioplene et al., 2015), which may be related to language learning ability or other factors that contribute to it. Published work on foreign language aptitude, however, has implicated cortical areas (for example Hu et al., 2013; Reiterer et al., 2011). Further, although all of our participants had obtained, or were engaged in study for, post-graduate degrees and thus relatively homogeneous in terms of their educational levels, we acknowledge that the data presented here did not incorporate information about socio-economic status, IQ or immigrant status, and that were therefore unable to control for these factors. In addition, it is possible that by looking only at individuals speaking more than two languages, our analysis was not sensitive to any structural changes that are related to the categorical leap from mono-to multilingual. This last point may also explain why the analysis revealed no relationship between LEXP and pallidal or thalamic morphology, in contrast to previous investigations (e.g. Burgaleta et al., 2016; Pliatsikas et al., 2016) that have reported a significant difference between monolingual and bilingual populations in these structures. In addition, Pliatsikas and colleagues (2016) report an effect of immersion on the monolingual vs bilingual subcortical differences that they find. The participants in the present study had highly variable levels of immersion in their non-native languages, and we were not able to assess the potential impact of immersion in third language (and beyond) environments, which may well have effects on subcortical brain structure.

These results are, nonetheless, consistent with the notion that the caudate nucleus is important in polyglot language control, and that multilingual expertise has consequences for structures implicated in a wide range of cognitive functions, including those associated with the bilingual advantage. We would argue that the data suggest that impact of multilingualism is not merely categorical but graded as a function multilingual experience. This represents an intriguing step forward in our understanding of the mechanisms of polyglot language control, which appear to exhibit ongoing plasticity in the face of increasing demands.

## Funding

This work was supported by the Swiss National Science Foundation Grants PP00P3_133701 and PP00P3_163756 awarded to NG.

## Acknowledgments

The authors are grateful to three anonymous reviewers whose helpful suggestions allowed us to substantially improve the manuscript. We also wish to express our gratitude to the staff at the Brain and Behaviour Lab at the University of Geneva and at the Lausanne University Medical Centre who supported the data acquisition.

## Conflict of Interest

The authors declare that they have no conflict of interest.

## Ethical approval

All procedures performed were in accordance with the standards of the local research ethics committees of the University Hospitals of Geneva and Lausanne, and with the 1964 Helsinki declaration and its later amendments.

## Footnotes

1 The following weights were used: 1) Proficiency: not fluent= 1, somewhat fluent = 2, moderately fluent=3, quite fluent=4, very fluent=5, native=6; 2) Age of acquisition: ages >21 years old = 1, ages 13-20 years old = 2, ages 7-12 years old=3, ages 1-6 years old=4, at birth=5.

2 Of these 67, 33 were multilingual individuals constituting a control group matched to an experimental sample of 34 trainee simultaneous interpreters. The trainees were scanned prior to the onset of their simultaneous interpretation training.

3 Although this might be considered a marginally-significant trend when applying a one-tailed test, we did not have a directional hypothesis on this question. We hope that future work will help to resolve whether putative language-mediated effects on left caudate nucleus structure are sensitive to age of acquisition of a second language.

## References

Abutalebi, J. (2008). Neural aspects of second language representation and language control. Acta psychologica, 128(3), 466–478.

Abutalebi, J., Annoni, J. M., Zimine, I., Pegna, A. J., Seghier, M. L., Lee-Jahnke, H., … Khateb, A. (2008). Language control and lexical competition in bilinguals: an event-related FMRI study. Cereb Cortex, 18(7), 1496–1505. doi:10.1093/cercor/bhm182

Abutalebi, J., Canini, M., Della Rosa, P. A., Sheung, L. P., Green, D. W., & Weekes, B. S. (2014). Bilingualism protects anterior temporal lobe integrity in aging. Neurobiol Aging, 35(9), 2126–2133. doi:10.1016/j.neurobiolaging.2014.03.010

Abutalebi, J., Della Rosa, P. A., Gonzaga, A. K., Keim, R., Costa, A., & Perani, D. (2013). The role of the left putamen in multilingual language production. Brain Lang, 125(3), 307–315. doi:10.1016/j.bandl.2012.03.009

Abutalebi, J., & Green*, D. W. (2008). Control mechanisms in bilingual language production: Neural evidence from language switching studies. Language and Cognitive Processes, 23(4), 557–582. doi:10.1080/01690960801920602

Abutalebi, J., Guidi, L., Borsa, V., Canini, M., Della Rosa, P. A., Parris, B. A., & Weekes, B. S. (2015). Bilingualism provides a neural reserve for aging populations. Neuropsychologia, 69, 201–210. doi:10.1016/j.neuropsychologia.2015.01.040

Agostinelli, C., & Library, S. L. A. T. E. C. C. M. (2015). wle: Weighted Likelihood Estimation. In.

Bak, T. H. (2016). The impact of bilingualism on cognitive ageing and dementia: Finding a path through a forest of confounding variables. Linguistic Approaches to Bilingualism, 6(1-2), 205–226. doi:10.1075/lab.15002.bak

Becker, T. M., Prat, C. S., & Stocco, A. (2016). A network-level analysis of cognitive flexibility reveals a differential influence of the anterior cingulate cortex in bilinguals versus monolinguals. Neuropsychologia, 85, 62–73. doi:10.1016/j.neuropsychologia.2016.01.020

Berken, J. A., Gracco, V. L., & Klein, D. (2017). Early bilingualism, language attainment, and brain development. Neuropsychologia, 98, 220–227. doi:10.1016/j.neuropsychologia.2016.08.031

Bialystok, E. (2011). Reshaping the mind: the benefits of bilingualism. Can J Exp Psychol, 65(4), 229–235. doi:10.1037/a0025406

Bialystok, E. (2017). The bilingual adaptation: How minds accommodate experience. Psychol Bull, 143(3), 233–262. doi:10.1037/bul0000099

Bialystok, E., Craik, F. I., & Luk, G. (2012). Bilingualism: consequences for mind and brain. Trends Cogn Sci, 16(4), 240–250. doi:10.1016/j.tics.2012.03.001

Burgaleta, M., Sanjuan, A., Ventura-Campos, N., Sebastian-Galles, N., & Avila, C. (2016). Bilingualism at the core of the brain. Structural differences between bilinguals and monolinguals revealed by subcortical shape analysis. Neuroimage, 125, 437–445. doi:10.1016/j.neuroimage.2015.09.073

Costumero, V., Rodriguez-Pujadas, A., Fuentes-Claramonte, P., & Avila, C. (2015). How Bilingualism Shapes the Functional Architecture of the Brain: A Study on Executive Control in Early Bilinguals and Monolinguals. Human brain mapping, 36(12), 5101–5112. doi:10.1002/hbm.22996

Crinion, J., Turner, R., Grogan, A., Hanakawa, T., Noppeney, U., Devlin, J. T., … Price, C. J. (2006). Language control in the bilingual brain. Science, 312(5779), 1537–1540. doi:10.1126/science.1127761

Cummine, J., & Boliek, C. A. (2013). Understanding white matter integrity stability for bilinguals on language status and reading performance. Brain Struct Funct, 218(2), 595–601. doi:10.1007/s00429-012-0466-6

Diamond, J. (2010). The Benefits of Multilingualism. Science, 330(6002), 332–333. doi:10.1126/science.1195067

Dijkstra, T., & Van Heuven, W. J. B. (2002). The architecture of the bilingual word recognition system: From identification to decision. Bilingualism: Language and Cognition, 5(03), 175–197.

García-Pentón, L., Fernández García, Y., Costello, B., Duñabeitia, J. A., & Carreiras, M. (2015). The neuroanatomy of bilingualism: how to turn a hazy view into the full picture. Lang Cogn Neurosci, 31(3), 303–327. doi:10.1080/23273798.2015.1068944

Garcia-Penton, L., Perez Fernandez, A., Iturria-Medina, Y., Gillon-Dowens, M., & Carreiras, M. (2014). Anatomical connectivity changes in the bilingual brain. Neuroimage, 84, 495–504. doi:10.1016/j.neuroimage.2013.08.064

Gold, B. T., Johnson, N. F., & Powell, D. K. (2013). Lifelong bilingualism contributes to cognitive reserve against white matter integrity declines in aging. Neuropsychologia, 51(13), 2841–2846. doi:10.1016/j.neuropsychologia.2013.09.037

Grahn, J. A., Parkinson, J. A., & Owen, A. M. (2008). The cognitive functions of the caudate nucleus. Prog Neurobiol, 86(3), 141–155. doi:10.1016/j.pneurobio.2008.09.004

Grazioplene, R. G., S, G. R., Gray, J. R., Rustichini, A., Jung, R. E., & DeYoung, C. G. (2015). Subcortical intelligence: caudate volume predicts IQ in healthy adults. Hum Brain Mapp, 36(4), 1407–1416. doi:10.1002/hbm.22710

Green, D. W., & Abutalebi, J. (2013). Language control in bilinguals: The adaptive control hypothesis. J Cogn Psychol, 25(5), 515–530. doi:10.1080/20445911.2013.796377

Grogan, A., Parker-Jones, O., Ali, N., Crinion, J., Orabona, S., Mechias, M. L., … Price, C. J. (2012). Structural correlates for lexical efficiency and number of languages in non-native speakers of English. Neuropsychologia, 50(7), 1347–1352. doi:10.1016/j.neuropsychologia.2012.02.019

Hämäläinen, S., Sairanen, V., Leminen, A., & Lehtonen, M. (2017). Bilingualism modulates the white matter structure of language-related pathways. Neuroimage, 152(6), 249–257.

Hervais-Adelman, A., Moser-Mercer, B., & Golestani, N. (2011). Executive control of language in the bilingual brain: integrating the evidence from neuroimaging to neuropsychology. Front Psychol, 2, 234. doi:10.3389/fpsyg.2011.00234

Hervais-Adelman, A., Moser-Mercer, B., Michel, C. M., & Golestani, N. (2015). fMRI of Simultaneous Interpretation Reveals the Neural Basis of Extreme Language Control. Cereb Cortex, 25(12), 4727–4739. doi:10.1093/cercor/bhu158

Hervais-Adelman, A., Moser-Mercer, B., Murray, M. M., & Golestani, N. (2017). Cortical thickness increases after simultaneous interpretation training. Neuropsychologia, 98, 212–219. doi:10.1016/j.neuropsychologia.2017.01.008

Higby, E., Kim, J., & Obler, L. K. (2013). Multilingualism and the Brain. Annual Review of Applied Linguistics, 33, 68–101. doi:10.1017/S0267190513000081

Hu, X., Ackermann, H., Martin, J. A., Erb, M., Winkler, S., & Reiterer, S. M. (2013). Language aptitude for pronunciation in advanced second language (L2) learners: behavioural predictors and neural substrates. Brain Lang, 127(3), 366–376. doi:10.1016/j.bandl.2012.11.006

Indefrey, P. (2006). A meta-analysis of hemodynamic studies on first and second language processing: Which suggested differences can we trust and what do they mean? Language Learning, 56, 279–304. doi:DOI 10.1111/j.1467-9922.2006.00365.x

Jenkinson, M., Beckmann, C. F., Behrens, T. E., Woolrich, M. W., & Smith, S. M. (2012). Fsl. Neuroimage, 62(2), 782–790. doi:10.1016/j.neuroimage.2011.09.015

Kroll, J. F., van Hell, J. G., Tokowicz, N., & Green, D. W. (2010). The Revised Hierarchical Model: A critical review and assessment. Biling (Camb Engl), 13(3), 373–381. doi:10.1017/S136672891000009X

Luk, G., Bialystok, E., Craik, F. I., & Grady, C. L. (2011). Lifelong bilingualism maintains white matter integrity in older adults. J Neurosci, 31(46), 16808–16813. doi:10.1523/JNEUROSCI.4563-11.2011

Luk, G., & Pliatsikas, C. (2016). Converging diversity to unity: commentary on The neuroanatomy of bilingualism. Language Cognition and Neuroscience, 31(3), 349–352. doi:10.1080/23273798.2015.1119289

Mechelli, A., Crinion, J. T., Noppeney, U., O’Doherty, J., Ashburner, J., Frackowiak, R. S., & Price, C. J. (2004). Neurolinguistics: structural plasticity in the bilingual brain. Nature, 431(7010), 757. doi:10.1038/431757a

Morgan-Short, K., Steinhauer, K., Sanz, C., & Ullman, M. T. (2012). Explicit and implicit second language training differentially affect the achievement of native-like brain activation patterns. Journal of Cognitive Neuroscience, 24%6(4), 933–947 %&. doi:10.1162/jocn_a_00119

Paap, K. R., & Greenberg, Z. I. (2013). There is no coherent evidence for a bilingual advantage in executive processing. Cogn Psychol, 66(2), 232–258. doi:10.1016/j.cogpsych.2012.12.002

Paap, K. R., Johnson, H. A., & Sawi, O. (2015). Bilingual advantages in executive functioning either do not exist or are restricted to very specific and undetermined circumstances. Cortex, 69, 265–278. doi:10.1016/j.cortex.2015.04.014

Patenaude, B., Smith, S. M., Kennedy, D. N., & Jenkinson, M. (2011). A Bayesian model of shape and appearance for subcortical brain segmentation. Neuroimage, 56(3), 907–922. doi:10.1016/j.neuroimage.2011.02.046

Pliatsikas, C., DeLuca, V., Moschopoulou, E., & Saddy, J. D. (2016). Immersive bilingualism reshapes the core of the brain. Brain Struct Funct. doi:10.1007/s00429-016-1307-9

Pliatsikas, C., Johnstone, T., & Marinis, T. (2014). Grey Matter Volume in the Cerebellum is Related to the Processing of Grammatical Rules in a Second Language: A Structural Voxel-based Morphometry Study. The Cerebellum, 13(1), 55–63. doi:10.1007/s12311-013-0515-6

Pliatsikas, C., Moschopoulou, E., & Saddy, J. D. (2015). The effects of bilingualism on the white matter structure of the brain. Proc Natl Acad Sci U S A, 112(5), 1334–1337. doi:10.1073/pnas.1414183112

R Core Team. (2015). R: A Language and Environment for Statistical Computing. Vienna, Austria: R Foundation for Statistical Computing. Retrieved from https://www.R-project.org/

Reiterer, S. M., Hu, X., Erb, M., Rota, G., Nardo, D., Grodd, W., … Ackermann, H. (2011). Individual differences in audio-vocal speech imitation aptitude in late bilinguals: functional neuroimaging and brain morphology. Front Psychol, 2, 271. doi:10.3389/fpsyg.2011.00271

Ressel, V., Pallier, C., Ventura-Campos, N., Diaz, B., Roessler, A., Avila, C., & Sebastian-Galles, N. (2012). An effect of bilingualism on the auditory cortex. J Neurosci, 32(47), 16597–16601. doi:10.1523/JNEUROSCI.1996-12.2012

Sebastian, R., Laird, A. R., & Kiran, S. (2011). Meta-analysis of the neural representation of first language and second language. Applied Psycholinguistics, 32(4), 799–819. doi:10.1017/S0142716411000075

Smith, S. M., & Nichols, T. E. (2009). Threshold-free cluster enhancement: addressing problems of smoothing, threshold dependence and localisation in cluster inference. Neuroimage, 44(1), 83–98. doi:10.1016/j.neuroimage.2008.03.061

Stein, M., Federspiel, A., Koenig, T., Wirth, M., Strik, W., Wiest, R., … Dierks, T. (2012). Structural plasticity in the language system related to increased second language proficiency. Cortex, 48(4), 458–465. doi:10.1016/j.cortex.2010.10.007

Stein, M., Winkler, C., Kaiser, A., & Dierks, T. (2014). Structural brain changes related to bilingualism: does immersion make a difference? Front Psychol, 5, 1116. doi:10.3389/fpsyg.2014.01116

Stocco, A., Lebiere, C., & Anderson, J. R. (2010). Conditional routing of information to the cortex: a model of the basal ganglia’s role in cognitive coordination. Psychol Rev, 117(2), 541–574. doi:10.1037/a0019077

Stocco, A., Yamasaki, B., Natalenko, R., & Prat, C. S. (2014). Bilingual brain training: A neurobiological framework of how bilingual experience improves executive function. International Journal of Bilingualism, 18(1), 67–92. doi:10.1177/1367006912456617

Wang, J., Zamar, R., Marazzi, A., Yohai, V., Salibian-Barrera, M., Maronna, R., … Konis., K. (2014). robust: Robust Library. Retrieved from https://CRAN.R-project.org/package=robust

Woumans, E., & Duyck, W. (2015). The bilingual advantage debate: Moving toward different methods for verifying its existence. Cortex, 73, 356–357. doi:10.1016/j.cortex.2015.07.012

Zatorre, R. J., Fields, R. D., & Johansen-Berg, H. (2012). Plasticity in gray and white: neuroimaging changes in brain structure during learning. Nat Neurosci, 15(4), 528–536. doi:10.1038/nn.3045

Zou, L., Ding, G., Abutalebi, J., Shu, H., & Peng, D. (2012). Structural plasticity of the left caudate in bimodal bilinguals. Cortex, 48(9), 1197–1206. doi:10.1016/j.cortex.2011.05.022

